# Rank-based deep learning from citizen-science data to model plant communities

**DOI:** 10.1101/2023.05.30.542843

**Authors:** Philipp Brun, Dirk N. Karger, Damaris Zurell, Patrice Descombes, Lucienne C. de Witte, Riccardo de Lutio, Jan Dirk Wegner, Niklaus E. Zimmermann

## Abstract

In the age of big data, scientific progress is fundamentally limited by our capacity to extract critical information. We show that recasting multispecies distribution modeling as a ranking problem allows analyzing ubiquitous citizen-science observations with unprecedented efficiency. Based on 6.7M observations, we jointly modeled the distributions of 2477 plant species and species aggregates across Switzerland, using deep neural networks (DNNs). Compared to commonly-used approaches, multispecies DNNs predicted species distributions and especially community composition more accurately. Moreover, their setup allowed investigating understudied aspects of ecology: including seasonal variations of observation probability explicitly allowed approximating flowering phenology, especially for small, herbaceous species; reweighting predictions to mirror cover-abundance allowed mapping potentially canopy-dominant tree species nationwide; and projecting DNNs into the future allowed assessing how distributions, phenology, and dominance may change. Given their skill and their versatility, multispecies DNNs can refine our understanding of the distribution of plants and well-sampled taxa in general.

## Main

During the last decade, we have witnessed a shift away from traditional expert-led biodiversity surveys towards data gathered via fast-growing citizen-science platforms^1, 2^. This development has resulted in an exponential increase in observational data, providing detailed information not only on locations of species observations, but also on their temporal dynamics^3, 4^. Such information is dearly needed to better understand biodiversity patterns and their underlying drivers, and to inform efficient conservation actions^5^. Yet, most citizen-science observations are made opportunistically which leads to significant spatial, temporal, and taxonomic biases^6^ in the data. Spatial biases are typically related to the accessibility of a location^7^, while temporal biases are linked to periods during which species are readily observed and/or easily identified (e.g. the flowering period). Taxonomic biases exist because citizen scientists prefer to report occurrences of conspicuous and charismatic species and taxa^8^. For analyzing and biodiversity^9^ these biases pose a major challenge, and therefore methods and approaches that can efficiently handle them are indispensable.

During recent decades, species distribution models (SDMs) have been among the most used tools to model biodiversity from species observations^10^. SDMs are a host of algorithms used to characterize habitat suitability and potential distributions of individual species as a function of their environmental niches^11^. When fitting SDMs with opportunistic citizen-science data, disentangling species’ environmental preferences from spatial sampling bias is a key challenge^12^. Sampling bias can be reduced by spatial or environmental thinning and pooling of observations^13^, and by manipulating the background points (or pseudoabsences)^14^. Thinning observations, however, leads to a considerable loss of information, which is then lacking to investigate aspects like the seasonal turnover of biodiversity. Moreover, sophisticated procedures of spatial bias correction are computationally demanding when applied to many species^15^.

Driven by their success in data science^16^, deep neural networks (DNNs) have become an increasingly popular alternative to model biodiversity from observational data^17–23^. Compared to SDMs, DNN-based modelling frameworks offer interesting new perspectives. They allow considering the spatial configuration of a landscape as predictor^21^, and they can cope with huge data sets^20, 21^. Additionally, DNN-based approaches typically model the distributions of many species jointly, which can be an efficient way of handling spatial biases. If observations of the modelled species depict similar sampling bias, the risk of confusing it with environmental preferences is reduced without the need to sacrifice observations through thinning. This, in turn, allows harnessing information from the full dataset, for example, the explicit consideration of seasonal effects on observed biodiversity^18, 19^. Immediate ways to cope with taxonomic reporting bias, on the other hand, are not offered by DNNs.

DNNs use cost functions to quantify how well model predictions match with observations. Hitherto, DNNs have often been optimized with cost functions that are not adequate for presence-only data. The standard cost function in multiclassification DNNs is the cross-entropy loss^24^ (CEL). This loss gets smaller the closer the predicted probabilities for absent classes are to zero, and the closer the probabilities for the present classes are to one. Like many common cost functions, the CEL therefore assumes absences for all but the reported species. This assumption is not met in citizen-science data that can contain large numbers of false absences, meaning that many species may be present but are not reported. A cost function that is more appropriate to citizen-science data should consider only presences and optimize the DNN to predict the observed species to be among the most relevant to occur, given environmental conditions. A similar criterion is used, for example, when training a search engine to rank the relevance of websites with respect to a given search query^25^. Although less common in ecology, many problems in data science require minimizing ranks, and various cost functions have been developed to address this problem^26^. Some of them, in particular the Normalized Discounted Cumulative Gain^27^ (NDCG), can readily be employed in multispecies DNNs^28^.

Here, we introduce rank-based deep learning to modeling the distributions of large numbers of species based on citizen-science data. Using occurrence observations of 2477 vascular plant species in Switzerland, we demonstrate that multispecies DNNs optimized with the NDCG cost function can accurately predict observation probabilities of thousands of species, both seasonally resolved and across large spatial scales. To do so, we jointly modeled all quality-filtered observations of the National Data and Information Center on the Swiss Flora (Info Flora, www.infoflora.ch), including 6.7 million citizen-science observations (Extended Data Fig. 1). We trained two versions of DNNs: a high-resolution (25×25 m) version, considering the full data and 18 environmental predictors, and, for a model comparison, a low-resolution (100×100 m) version, considering 4.6 million observations and ten environmental predictors. At both resolutions we complemented environmental predictors with two seasonal predictors based on a sine-cosine mapping of day of year^19^, in order to also model observation probability in response to time (season). We compared the performance of low-resolution DNNs trained with NDCG and CEL cost functions to the performance of stacked predictions of SDMs (SSDMs) that were trained with the same coarse-resolution environmental but without seasonal predictors. Moreover, to highlight the insight that can be derived from multispecies DNNs, we focused on three aspects that are difficult to analyze with traditional approaches: (1) mapping variations in the phenology of selected forb species; (2) mapping potentially dominant tree species; and (3) projecting spatiotemporal distributions of graminoid species under severe climate change.

## Results

### Performance

DNN fits trained with the NDCG cost function made better predictions to left-out citizen science observations than DNN fits trained the CEL cost function and SSDMs. We measured predictive performance based on the ranks of observed species in model predictions for 12’187 taxonomically balanced test observations, with ranks close to one representing highest performance and ranks close to the maximum of 2477 species representing lowest performance. We weighted the predicted ranks with the number of training observations per species to obtain scores that represent typical field observations, and we used paired Wilcoxon tests and the Holm correction^29^ for multiple comparisons to test for significant differences between models. Observed species had significantly lower ranks in predictions of the low-resolution NDCG DNN than in predictions of the low-resolution CEL DNN and SSDMs, with weighted medians of 74, 76, and 175, respectively (Fig. 1a; *p* ≤ 0.001 for all pairwise comparisons; *n* = 12’187). Training the NDCG DNN with the high-resolution data set further improved predictive performance, lowering the weighted median to 43. Training the low-resolution NDCG DNN without seasonal predictors, i.e., with the same information as SSDMs, still yielded distinctly higher performance, with a median weighted rank of 87 (Extended Data Fig. 2). The relative advantage of NDCG DNNs was even higher when measured with top-1 and top-5 accuracy (Extended Data Fig. 2).

**Fig. 1.**
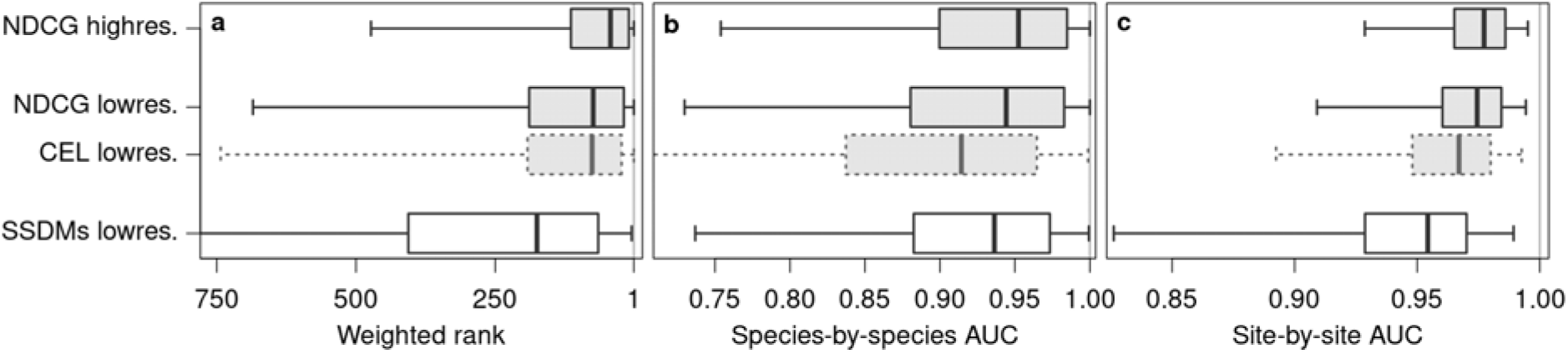
Performance of the deep neural networks (DNNs) trained with different data sets and cost functions in comparison to stacked species distribution models (SSDMs). **a**, weighted ranks inferred from a left-out test set of 12’187 citizen-science observations. **b**, species-by-species AUC measuring the performance of predicting the distributions of 1339 individual species based on independent survey data. **c**, site-by-site AUC measuring the performance of predicting the composition communities at 1489 sites based on independent survey data. Boxes of DNNs are shown in gray and boxes of SSDMs are shown in white. Results of the DNN trained with cross-entropy loss (CEL) are shown with dashed lines; results of DNNs trained with the normalized discounted cumulative gain (NDCG) are shown with solid lines. Central lines in the boxplots indicate medians, boxes indicate interquartile ranges, and whiskers indicate 2.5 and 97.5 percentiles.

NDCG DNNs also showed a higher capacity than SSDMs to predict individual species’ distributions and in particular community composition. We used 1489 regularly distributed plant community inventories from the Swiss Biodiversity Monitoring program (www.biodiversitymonitoring.ch) for independent validation. Annual summaries (see methods) of low-resolution NDCG DNN predictions showed a slightly but significantly higher performance in predicting species distributions than SSDMs (Fig. 1b; *p* ≤ 0.001 for all comparisons; *n* = 1339), with a median species-by-species area under the curve^30^ (AUC) of 0.944 versus 0.936. The advantage of NDCG DNNs was even higher when predicting community composition. Median site-by-site AUC was 0.954, 0.974, and 0.977, for SSDMs, the low-resolution NDCG DNN, and the high-resolution NDCG DNN, respectively (Fig. 1c; *p* ≤ 0.001 for all comparisons; *n* = 1489). CEL DNNs also performed significantly better than SSDMs in predicting community composition, but significantly worse in predicting species distributions (Fig. 1b,c).

### Predicting spatial variations in species’ phenology

Plant phenology is the study of seasonally recurring phenomena in plant development, which are important indicators of ecosystem functions and their susceptibility to climate change^31^. Despite increasing attention, our capacity of modelling phenological events is still limited^31, 32^. We explored the spatial patterns in the seasonal variation of observation probability and their links to phenology for a selected set of common forb species by mapping the timing of peak observation probability (*t_p_max_*).

At the scale of Switzerland *t_p_max_* showed distinct spatial patterns for the evaluated species. *t_p_max_* of bugle (*Ajuga reptans*) started in April at lower elevations, while at higher elevations in the Prealps and the Jura mountains it was around early June (Fig. 2a). The case was similar for common twayblade (*Listera ovata*) that showed an even higher plasticity in *t_p_max_* (Fig. 2b). Also, for the common dandelion (*Taraxacum officinale* aggr.) *t_p_max_* was quite variable (Fig. 2c). Yet, in some regions of the Swiss plateau, the pattern was more scattered than it was for the former two species. The seasonal progression of observation probability at a few selected locations indicates that this scattering results from double peaks in observation probability: on the Swiss plateau observation probability peaked twice, once around the end of April and once in August. This phenology was distinct from the phenology in the Jura mountains or in the high-elevation valleys of the eastern alps, where *t_p_max_* was between end of May and June, but also form the phenology in the lowland areas of the central valley and southern Ticino, where *t_p_max_* was in early April without a second peak later in the season.

**Fig. 2.**
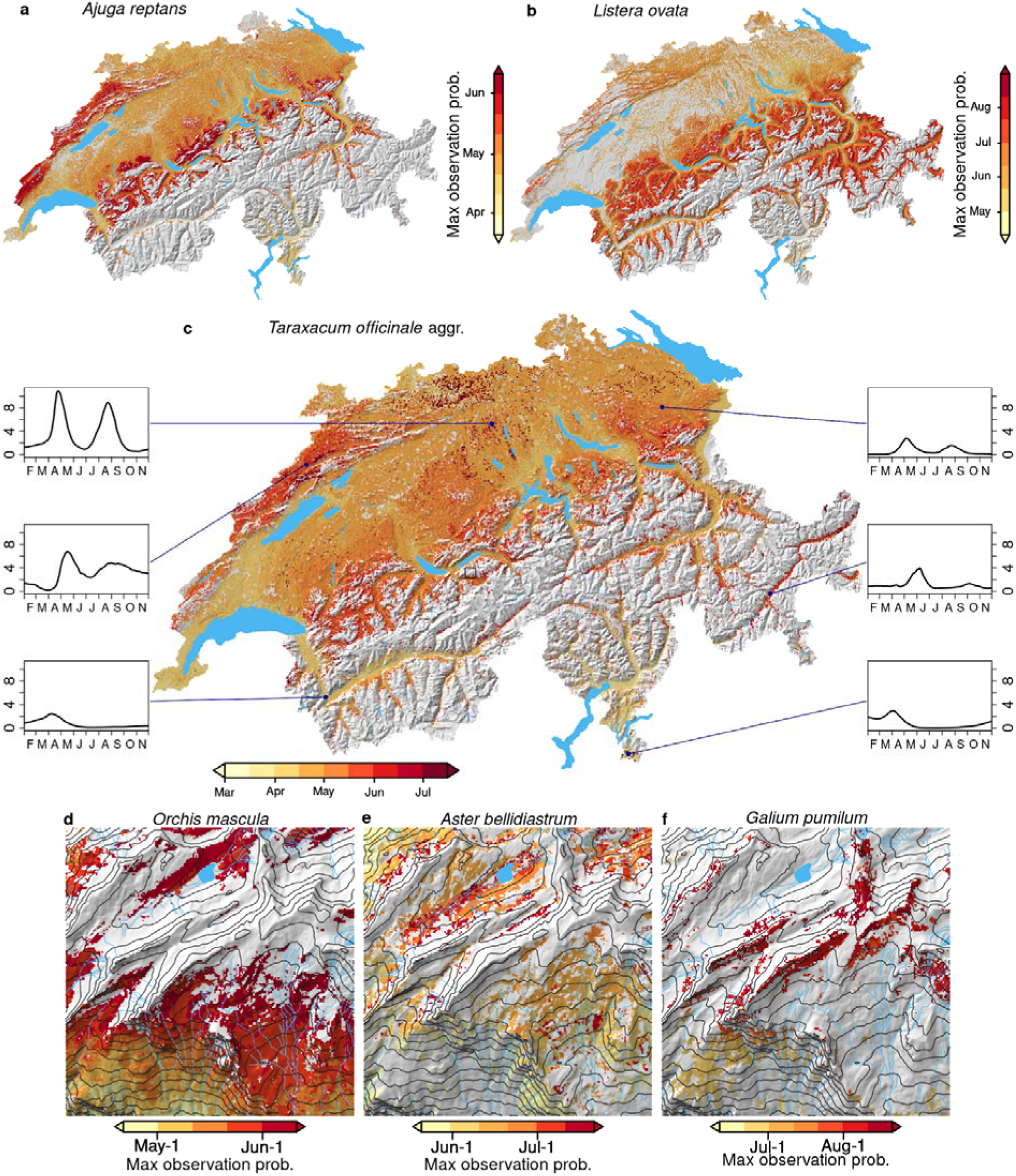
Seasonal maxima of observation probability for selected species. Panels **a**-**c** show the timing of highest observation probability across the distribution ranges of *Ajuga reptans*, *Listera ovata* and *Taraxacum officinale* aggr., respectively. Shown are pixels with a maximum daily observation probability ≥0.01 between March and September. Insets in panel **c** illustrate the seasonal evolution of observation probability (in percent, from February to November) at six selected locations within the distribution range of *T. officinale* aggr. Panels **d**-**f** also show timing of peak observation probability, but for a 5×5 km square in the region of Faulhorn (outlined on the map in panel **c**) for *Orchis mascula*, *Aster bellidiastrum*, and *Galium pumilum* aggr., respectively.

*t_p_max_* also showed striking local patterns in a mountainous 5×5 km square north of Grindelwald. For early-purple orchid (*Orchis mascula,* Fig. 2d), false aster (*Aster bellidiastrum,* Fig. 2e), and slender bedstraw (*Galium pumilum* aggr., Fig. 2f) *t_p_max_* delayed rapidly with elevation. To a lesser degree *t_p_max_* also responded to other terrain properties, such as slope and exposition, as in the case of early-purple orchid, or troughs, as in the case of false aster, affected *t_p_max_*.

Predictions of *t_p_max_* matched well with dates on which the selected species were seen blooming. We subsampled Info Flora observations for which phenology information was available and classified as ‘full bloom’ and compared their dates to corresponding predictions of *t_p_max_*. For the six selected species of forbs, median difference between observation dates and *t_p_max_* ranged from −2 to 10, while Spearman rank correlation ranged from 0.40 to 0.75 (Table 1).

**Table 1.** **Predictive performance of the timing of blooming for the species shown in Fig. 2.** *n* refers to number of test observations; *r* refers to correlation coefficient.

| Species | $n$ | Spearman $r$ | Bias (days) |
| --- | --- | --- | --- |
| *Ajuga reptans* | 649 | 0.56 | 4 |
| *Aster bellidiastrum* | 304 | 0.66 | 3.5 |
| *Galium pumilum* aggr. | 275 | 0.57 | 0 |
| *Listera ovata* | 9076 | 0.75 | 4 |
| *Orchis mascula* | 8415 | 0.50 | 10 |
| *Taraxacum officinale* aggr. | 656 | 0.40 | -2 |

### Predicting potentially dominant species

For many problems in ecology, ecosystem management, and conservation knowledge is necessary not only on species’ habitat suitability but also on potential dominance, especially for structurally important species like trees^33^. We estimated potential dominance for 37 tree species form high-resolution NDCG DNN predictions by reweighting observation probabilities to represent Braun-Blanquet^34^ cover-abundance scores in the canopy. This reweighting enforced species-specific sums of predicted probabilities across 12’911 sites of the Swiss forest vegetation database^35^ to be equal to sums of observed Braun-Blanquet scores, which also corrected for taxonomic reporting bias.

Across Switzerland, Norway spruce (*Picea abies*), especially in the Alps, and European beech (*Fagus sylvatica*), especially on the Swiss Plateau, were predicted to potentially dominate most often, in 43.5% and 32.0% of the wooded area, respectively (Fig. 3a). Other species were predicted to potentially dominate in a narrower range of environmental conditions, for example European larch (*Larix decidua,* 4.8%) and Swiss stone pine (*Pinus cembra*, 3.9%) in the alps, Scots pine (*Pinus sylvestris*, 5.1%) in the Central valley and the upper Rhine valley, and sweet chestnut (*Castanea sativa*, 1.7%) on low-elevation slopes in Ticino.

**Fig. 3.**
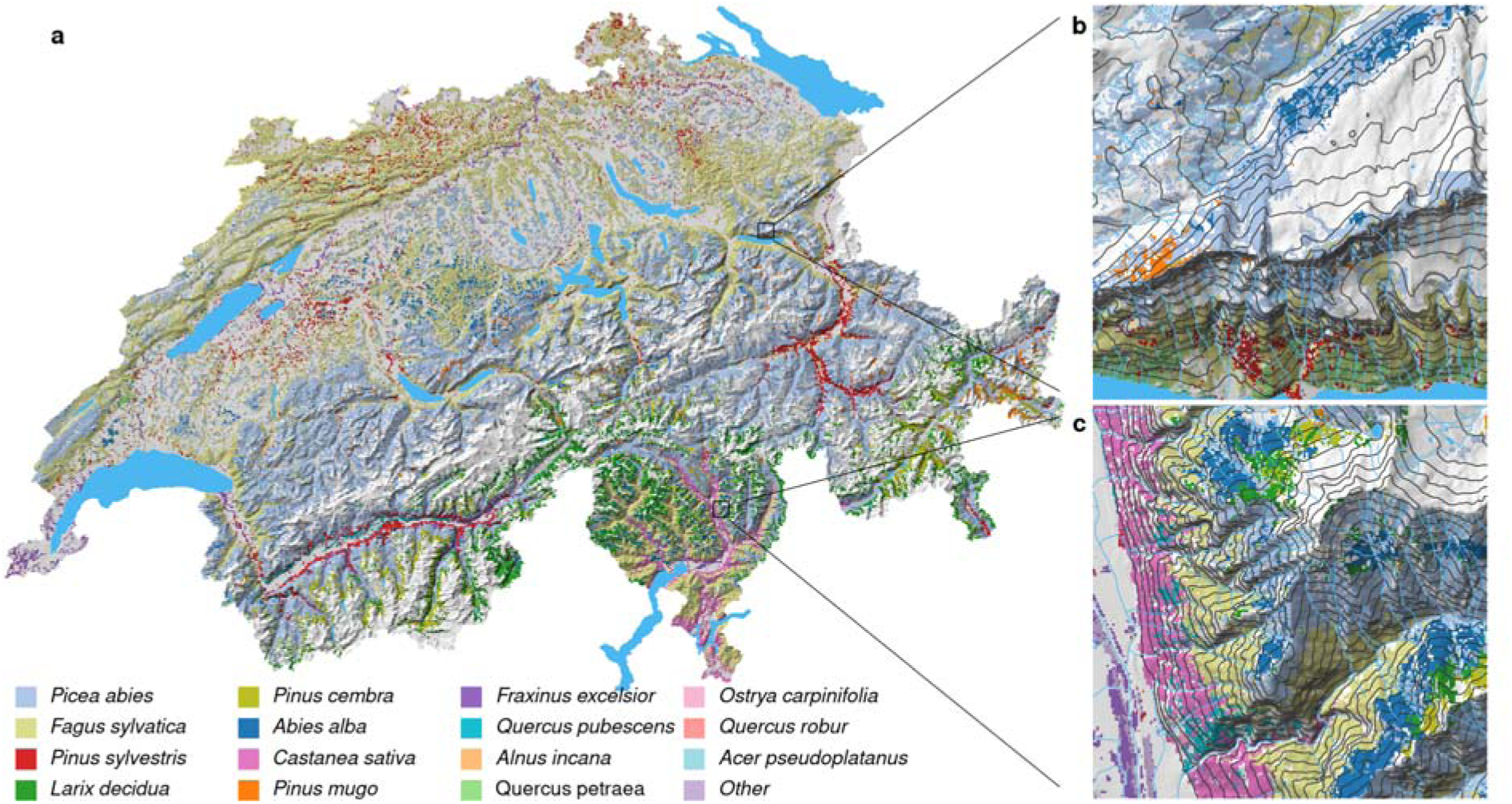
Potentially dominant canopy-forming tree species in wooded areas of Switzerland. **a**, for the entire country; **b**, for a selected 5×5 km square near Quinten; **c**, for a selected 5×5 km square near Biasca. Colors represent species with the highest weighted observation probability averaged from February to November (as indicated in the legend at the figure bottom). We compared observation probabilities of 37 tree species, as distinguished by the Swiss forest vegetation database^35^, and masked pixels with land cover classes without trees (see methods).

At the local scale, a turnover in potentially dominant tree species was predicted along gradients of elevation and exposition. On the steep, south-facing slope north of Walensee, sessile oak (*Quercus petraea*) was predicted to potentially dominate close to the lakeshore, while at higher elevations European beech and Scots pine took over (Fig. 3b). Norway spruce was predicted to take over potential dominance at relatively high elevations but it reached further downhill on the north-facing side, behind the ridge. On the eastern side of the Riviera valley, potentially dominant tree species showed a distinct horizontal layering (Fig. 3c). On the valley bottom, European ash (*Fraxinus excelsior*) and other species potentially dominated; at the lower slopes it was sweet chestnut intermixed with downy oak (*Quercus pubescens*); higher up European beech took over for an elevation band of several hundred meters before being displaced by Norway spruce and silver fir (*Abies alba*).

Predictions of potentially dominant species showed a reasonable agreement with observations from the National Forest Inventory^36^. Top-1 accuracy was 0.52 across 1867 regularly distributed sites evaluated. F_1_ scores were comparably high for common species like Norway spruce and European beech, but also for rarer alpine species like Swiss stone pine (Table 2). Predictions were less successful for widespread lowland species that rarely dominate large stands, such as European ash and sessile oak.

**Table 2.**
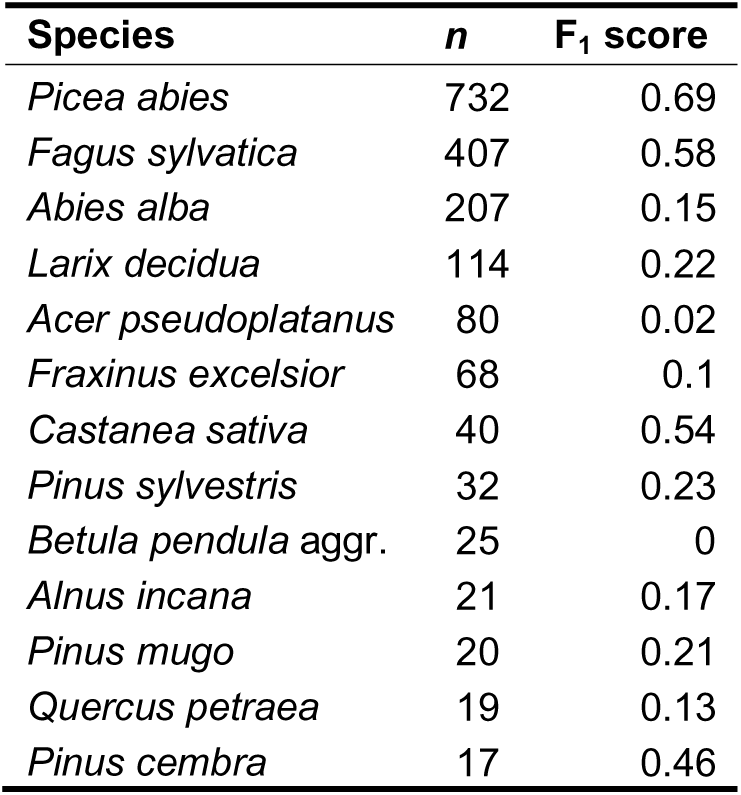
**Predictive performance for the 13 most-commonly dominating canopy tree species according to the National Forest Inventory.** *n* refers to number of test observations; F_1_ score is the harmonic mean of precision and recall.

| Species | *n* | $F_1$ score |
| --- | --- | --- |
| *Picea abies* | 732 | 0.69 |
| *Fagus sylvatica* | 407 | 0.58 |
| *Abies alba* | 207 | 0.15 |
| *Larix decidua* | 114 | 0.22 |
| *Acer pseudoplatanus* | 80 | 0.02 |
| *Fraxinus excelsior* | 68 | 0.1 |
| *Castanea sativa* | 40 | 0.54 |
| *Pinus sylvestris* | 32 | 0.23 |
| *Betula pendula* aggr. | 25 | 0 |
| *Alnus incana* | 21 | 0.17 |
| *Pinus mugo* | 20 | 0.21 |
| *Quercus petraea* | 19 | 0.13 |
| *Pinus cembra* | 17 | 0.46 |

### Predicting climate-change impact

Projections of phenology and potential dominance can also enrich assessments of climate change impact, which may allow for better-informed mitigation strategies. Using the low-resolution NDCG DNN, we predicted potential dominance of graminoid species along an elevational gradient (400-2900 m asl.) in the canton of Vaud under current conditions and under conditions expected in 2071-2100, assuming RCP8.5^37^. We reweighted predictions as for the tree species but based on 23’919 community plots from the dry meadows and pastures initiative^38^, and we sub-selected the twelve most-frequent potential dominators.

In May and June, under current conditions, the potentially most dominant species were predicted to be bulbous oat grass (*Arrhenatherum elatius*), erect brome (*Bromus errectus*) and blue moor-grass (*Sesleria caerula*) for the valley bottom, the lower slopes and the upper slopes, respectively (Fig. 4a, b). In July and August, erect brome still potentially dominated, while in the valley it was intermediate wheatgrass (*Elymus hispidus*) and in the upper slopes several species were predicted to partially dominate, including red fescue (*Festuca rubra* aggr.) at intermediate elevations, and purple fescue (*Festuca violacea* aggr.) at high elevations (Fig. 4c, d).

**Fig. 4.**
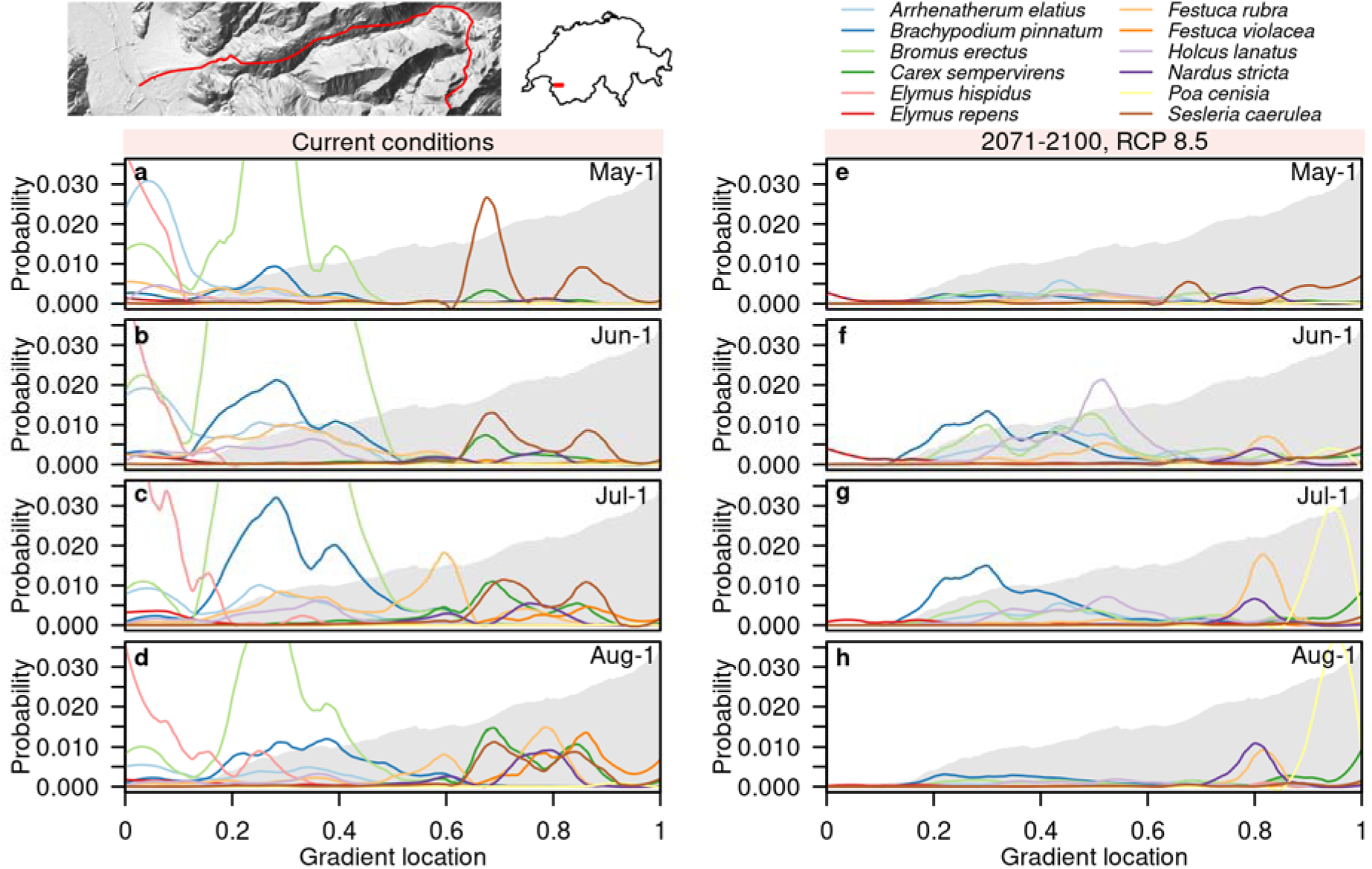
Reweighted observation probabilities of graminoid taxa along an elevational gradient in the canton of Vaud (see inset maps). Panels **a**-**d** illustrate bias-corrected observation probabilities along the gradient for the twelve most likely observed graminoid species (see legend) under current conditions for May-1, June-1, July-1, and August-1, respectively. Panels **e**-**h** illustrate equivalent observation probabilities for the period 2071-2100, assuming the emission scenario RCP8.5. Probabilities along the gradient are represented as local polynomial regression fits (see methods). Gray shading in the background represents the elevation profile ranging from 400 to 2900 m asl. For an SSDM-based version see Extended Data Fig. 3.

For the period 2071-2100, potential dominance was projected to change, and species to shift their ranges (defined as gravity centers of dominance probability) towards higher elevations and to advance *t_p_max_* (Fig. 4e-h). In the valley and on the lower slopes couch grass (*Elymus repens*) and tor-grass (*Brachypodium pinnatum*), respectively, were projected to mostly take over potential dominance. Blue moor-grass and red fescue were projected to potentially dominate at higher elevations than under current conditions, and observation probabilities of erect brome relatively declined in July and August, with few remaining locations of potential dominance, mainly in May and June.

## Discussion

Our results highlight that when equipped with an appropriate cost function DNNs can efficiently learn the distributions of thousands of species based on millions of citizen-science observations, without the need of sacrificing observations to spatial bias correction. NDCG DNNs were better than SSDMs in all comparisons, but especially in ranking observed species and in predicting community composition, which are both multispecies problems. More importantly even, multispecies DNNs offer new and promising perspectives to ecological research.

Firstly, they can resolve spatial variations in phenology. For common forb species we have identified the timing of peak observation probability within pixels and revealed striking spatial patterns in detail. Our results suggested interesting phenomena, such as double peaks in the blooming of *T. officinale*, which have also been observed in other regions^39^. The correspondence of predicted *t_p_max_* with observations was satisfying, given the extended duration of flowering periods, their substantial interannual variations^40^, and the indirect nature of our approach. Future efforts may further improve estimates of *t_p_max_*by filtering training observations for blooming individuals as identified from auxiliary images^41^, and by using annual climate data. Understanding the spatial variations in species’ phenology can improve, for example, our understanding of speciation processes through reproductive isolation^42^, and the impact of phenological mismatches on the spatial variation of plant-pollinator networks^43^.

Secondly, multispecies DNNs can jointly predict observation probabilities for many species. We have shown that these observation probabilities may be reweighted to correct for taxonomic reporting bias and to account for cover-abundance. This way, we have generated a nationwide map of potentially dominant canopy-forming trees that is more detailed than existing alternatives^44, 45^ and that corresponds reasonably well to observations, given the relatively crude input on vegetation structure and that 45% of Swiss forests are not in a natural state^46^. Such a map cannot be created by stacking predictions from individual species’ models (Extended Data Fig. 4), and it would likely be less detailed without ubiquitous citizen-science observations. Predictive performance may improve if rank-based DNNs were confined to the relevant species and trained with detailed remote-sensing predictors^20, 33^. Maps of potentially dominant species may be useful for forest rangers to identify locally competitive species^47^, but comparable observation probabilities are also advantageous in other contexts. They can, for example, be used to quality control citizen-science observations or to improve image-based plant species identification^18, 19^.

The third perspective emerges from projecting climate-change impact on phenology and potential dominance. In addition to moving towards higher elevations, our DNN projected, e.g., erect brome to advance *t_p_max_*in the season, which matches well with the general expectation of advancing phenology in European grasslands^48^. Modeling climate-change impact on phenology at the species level allows quantifying to which extent plants can cope by shifting their growing season, which may improve their resilience^49^, and it allows assessing where phenological mismatches of interacting species may arise, for example between plants and pollinators^50^. Moreover, when potential dominance can be estimated, multispecies DNNs allow projecting how it may change, as we have shown for tor-grass and erect brome at intermediate elevations. And when such turnovers affect structurally important species, consequences at the ecosystem level may be dire. Accounting for phenology and potential dominance may critically refine climate-change impact assessments that so far have often focused on range shifts^51^.

Every year, citizen scientists report millions of species observations which are urgently needed to cope with the ongoing biodiversity crisis^5^ but are also biased and challenging to analyze. Rank-based multispecies DNNs are not only capable of skillfully modeling such data, they also offer unprecedented insight into the spatial patterns of species’ phenology and potential dominance, which allows comprehensive prognoses of climate-change impact. Sufficiently high observation numbers are a mandatory prerequisite to successfully build DNNs, which is why they may currently be most advantageous in well-sampled regions. But given that citizen-science contributions to biodiversity monitoring keep growing^2^, and that we have only started exploring this method, chances are high that it will become a standard item in the toolbox of tomorrows macroecologists.

## Material and Methods (3161)

### Overview

We trained different versions of multispecies DNNs, assessed their performance, and explored their seasonal and spatial predictions under current and future climate conditions. Different versions of low-resolution (100×100 m) multispecies DNNs were trained, using two cost functions and several predictor sets, and their performance was compared to stacked species distribution models that were fitted at the same low resolution. Additionally, we trained and evaluated high-resolution (25×25 m) DNNs and used them to map the timing of seasonal maxima in observation probability (*t_p_max_*), as well as potentially dominant trees at different spatial scales. Finally, using low-resolution DNNs, we investigated how predicted observation probabilities may be altered by the end of the century, assuming rapid climate change.

### Data

#### Observations

We worked with five sets of observational data: a main set of opportunistic citizen-science observations for model training, and four sets of expert-based survey data for independent validation and correction of taxonomic reporting bias. The opportunistic data set was provided by the National Data and Information Center on the Swiss Flora (Info Flora, www.infoflora.ch) and contained plant observations in Switzerland that were primarily made by citizen scientists. Observations were made during recent decades, with continuously increasing numbers per year and 80% of contributions after 1998 (Extended Data Fig. 5). We filtered these data for observations made in 1971 or more recently, for taxon labels at the level of species aggregates or finer, for identifications with high confidence as indicated by the users, for coordinates within Switzerland that are plausible for the species (as assessed by Info Flora), and for a coordinate uncertainty of no more than 100 meters. Then, we removed duplicates and pooled observations with subspecies-level labels with those of the corresponding species, and observations with species-level labels with those of the corresponding species aggregates (for the 12% of species that are part of species aggregates). For simplicity, we jointly referred to species and species aggregates as ‘species’ throughout the main text. We used two versions of Info Flora data: for low-resolution DNNs and for stacked SDMs we worked with an extraction that was made in April 2019, consisting of 5.1 million records after filtering. For stacked SDMs, we used this data set directly, but for multispecies DNNs we further filtered it for the availability of an observation date. In this step, we also excluded observations with time stamps representing the last or the first second of the year, which originated from observation time information with annual resolution. Filtering for observation date reduced the data set to 4.6 M observations. For the high-resolution analysis, we worked with a more recent extraction (November 2022) and applied a stricter 25-m-threshold to coordinate uncertainty and the same filter for observation date, resulting in 6.7 M filtered records.

We partitioned the Info Flora observations into training and test data and filtered for taxa with a minimum number of occurrences (Extended Data Fig. 1). Test data consisted of five randomly sampled observations per taxon with observation date and no more than 25 m coordinate uncertainty (three taxa were represented with four test observations). We sampled these test observations from the more recent version of the data set, wherever possible with observation dates after April 2019 (99.8% of cases) and where not possible with older observation dates. Training data of both versions of the opportunistic data set were then defined as all observations that were not part of the test set, except in case of stacked species distribution models, where we used all observations of the older version of the data set as training data. In addition to five test observations, we expected at least 20 training observations in both versions of the data set for a taxon to be considered, which was the case for 2477 species and species aggregates.

Surveys from the forest database and the dry meadows and pastures initiative were used to correct for taxonomic reporting bias. The Swiss forest vegetation database^35^ reported the cover-abundance of 41 tree and shrub species at three vertical layers for 14’860 survey sites within Switzerland and in neighboring regions. We filtered the data for surveys within Switzerland with full coverage of environmental data (see below) and summed species-level information to species aggregates, where applicable. Semi-quantitative cover-abundance information from the canopy-layer was translated from the original Braun-Blanquet scheme^34^ to percentages, following ref. ^52^. After filtering and aggregation, cover-abundance information was available for 37 taxa at 12’911 survey sites (Extended Data Fig. 1). The dry meadows and pastures initiative^38^ was run by the Swiss Federal Office for Environment and consists of cover-abundance information for almost 24’000 surveyed grassland communities. We focused on 91 graminoid taxa that are dominant and influence the appearance of at least one grassland, wetland or cropland habitat type in Switzerland, according to the TypoCH vegetation classification^53^, and conducted the same aggregation and filtering steps as for the surveys from the Swiss forest vegetation database. After processing, cover-abundance information was available for 23’919 surveys (Extended Data Fig. 1).

Expert-based surveys from the Swiss Biodiversity Monitoring Program^54^ (BDM, www.biodiversitymonitoring.ch), and from the National Forest Inventory^36^ (NFI, www.lfi.ch) were used for independent validation. Survey data from the BDM program were used to validate predicted distributions of individual species and community composition. We considered the BDM indicator Z9 (species diversity in habitats), containing 1562 sites that are regularly distributed across Switzerland. Between 2000 and 2021 each of these 10 m^2^ areas was revisited three to four times and all plant taxa present were recorded. Taxa were mostly identified at the species level, but identifications at the level of species aggregates were also common. We considered a taxon present at a site if it was observed in at least one revisitation. Where applicable, we pooled species to aggregates, yielding 1489 survey sites (Extended Data Fig. 1) with at least one species or species aggregate observed and full environmental data coverage (see below), and 1345 taxa observed within the survey sites. Survey data from the NFI were used to validate predicted potentially dominant tree species. The NFI samples forests on a regular grid with 1.4 km spacing (where forests occur, see Extended Data Fig. 1). The sampling procedure includes the species-level identification of all trees with ≥12 cm stem diameter at breast height in an inner circle covering 200 m^2^, and the identification of all trees with ≥36 cm diameter at breast height within an outer circle covering 500 m^2^. We considered survivor trees from the third (2004-2006) to the fourth (2009-2017) assessment^36^ and defined the species with most individuals with ≥12 cm stem diameter at breast height to be dominant. After filtering and pooling for the same 37 species and species aggregates as distinguished by the forest database, information on dominant tree taxa at 1867 sites was available.

#### Environmental predictors

We constructed a low-resolution and a high-resolution set of environmental predictors, representing vegetation structure, climate, soil conditions, topography, and season. Two separate sets were necessary to allow for a fair comparison between multispecies DNNs and stacked SDMs (which were trained in 2019), and to include the best data available for the high-resolution analysis. The low-resolution (100×100 m) set included 24 environmental variables, projected in the CH1903/LV03 projection system, and two predictors representing season. The low-resolution environmental layers originated from two recent studies investigating the distribution of Swiss plant species^15, 55^. The predictor group representing vegetation structure included canopy height, inner forest density, and mean and standard deviation of the normalized difference vegetation index (NDVI). Canopy height and inner forest density were estimated by the 25^th^ and 95^th^ height percentiles of normalized LiDAR returns above 40 cm^15, 56^. Mean and standard deviation of NDVI were calculated from Landsat measurements (https://espa.cr.usgs.gov/) taken during summer months (July to mid-September) between 2007 and 2015. Climate variables describe temperature, precipitation, and solar radiation (direct and diffuse) and were downscaled from reanalysis data^57^ to the EarthEnv-DEM90 digital elevation model^58^, using the CHELSA approach^59^. For temperature and precipitation, we considered averages and sums, respectively, for June to August, December to February, and for the whole year. For solar radiation, we only considered annual averages. In addition, we included CHELSA-based estimates of frost change frequency and growing degree days (with 5 °C baseline temperature) and plant indicator value-based estimates of continentality and light availability^55^ as climatic predictors. Soil conditions are represented by plant indicator value-based estimates of pH, moisture, moisture variability, aeration, humus, and nutrients^55^. Topography variables included terrain wetness index, terrain ruggedness index, and terrain position index, and were derived from the EarthEnv-DEM90 digital elevation model^58^. More details on the generation of these predictor variables are provided in refs ^55^ and ^15^. Finally, we used sine-cosine mapping to encode the season of observation (day of year) into two circular variables^19^, one of them roughly indicating day length and the other change in day length per day. Collinearity in this set of predictors overall was rather low, except for some predictors related to temperature and, to a lesser degree, precipitation (Extended Data Fig. 6). In the subsequent analyses predictors were subsampled in order to avoid pairs with high correlations (see below).

The high-resolution (25×25 m) predictor set roughly covered the same predictor groups but included some additional variables. High-resolution variables were compiled in the CH1903/LV95 projection system. For vegetation structure, we aggregated the country-wide vegetation height model with 1 m original horizontal resolution^60^ to estimate maximum and 25^th^ percentile of vegetation height in each 25×25 m cell. In addition, we calculated annual medians and seasonal differences in enhanced vegetation index (EVI) for the period 2018-2021. EVI data was calculated based on 10×10 m Sentinel 2 measurements^61^ and compiled to analysis-ready annual and seasonal medians by the Swiss Data Cube^62–66^. We removed values outside the valid range (−1 to 1), and reprojected and aggregated the layers to 25×25 m by average. Then, we aggregated annual, summer, and winter medians inter-annually by median and calculated the difference between summer and winter median. For a handful of pixels, no valid EVI measurements were available, so we approximated them with bilinear interpolation. We also considered data on canopy mixture (deciduous versus evergreen) in forested areas at 25×25 m original resolution^45^. We encoded this information as two variables, one representing deciduous forest cover and one representing evergreen forest cover, setting the values for unforested areas to zero. Related to vegetation structure, we also included information on land cover at 25×25 m original resolution^67^, encoding the 62 classes distinguished individually as binary factors (one hot encoding). High-resolution temperature and precipitation data were derived from the CHclim25 data set^68^. These data were derived from topography-dependent interpolation of station measurements to 25 m resolution. From the monthly climatologies for the period 1981-2010 we derived annual means and sums, for temperature and precipitation respectively, as well as annual ranges. Moreover, we calculated precipitation sums for the summer months (June to August). Terrain variables were derived by aggregating the swissAlti3D model, distributed by the Swiss Federal Office of Topography, from the original 2 m resolution to 25 m by average. From the elevation data (not used as predictor) we calculated aspect as well as the SAGA wetness index (using SAGA GIS^69^). In addition, we calculated the terrain ruggedness index for each 25×25 m cell (maximum minus minimum elevation across the covered 2×2 m cells). To represent solar radiation, soil conditions, and season, we used the same variables as in the low-resolution set, downscaling them, where applicable, with bilinear interpolation. Collinearity in the high-resolution predictor set was low (Extended Data Fig. 7), except for the association of mean annual temperature with annual temperature range (*r* = 0.74) and the association between annual precipitation and summer precipitation (*r* = 0.89). Unless otherwise noted, data were prepared in the R environment^70^ (R version 3.6.3) using the package terra^71^.

### Model configurations

#### Multispecies deep neural networks

##### Network architecture and training settings

We trained multispecies deep neural networks at low (100×100 m) and high (25×25 m) resolution. All DNNs were set up with six fully connected hidden layers and 380 nodes per layer, and the main predictor set at both resolutions included environmental and seasonal predictors. For low-resolution DNNs, we sub-selected ten environmental predictors from the low-resolution set. The primary criteria thereby were variable importance in species distribution models, and absolute Pearson correlation coefficients of no more than 0.7 (Extended Data Fig. 6 and Extended Data Table 1). Yet, we made one exception by choosing summer temperature over solar radiation, despite somewhat lower variable importance, to be able to better account for climate change effects. For the high-resolution DNN, we included the whole set of environmental predictors, as (1) only two predictor pairs with absolute Pearson correlation coefficients >0.7 existed (Extended Data Fig. 7), (2) our study area was heavily sampled, and (3) we did not use the high-resolution DNN for future projections. All predictor variables were rescaled to range from −1 to 1 prior to analysis. For low-resolution DNNs, we conducted a sensitivity analyses with an alternative predictor set consisting of environmental predictors only (no seasonal predictors. We trained all models first for 50 epochs with equal sampling probabilities of training observations for each taxon (by upsampling rare species), and then for 100 and 50 additional epochs with unmodified sampling probabilities for low-resolution and high-resolution DNNs, respectively. We chose a learning rate of 5 * 10^-4^ and 2.5 * 10^-4^, for low-resolution and high-resolution DNNs, respectively, which was reduced when the loss was plateauing, and optimized with the stochastic gradient descent method^72^, setting the momentum to 0.9. For the second part of training without weighting, initial learning rates were halved. Choosing a lower learning rate and less epochs for the high-resolution DNN prevented the cost function from reaching full saturation during training. This set-up led to minimally lower performance as evaluated by the Info Flora and BDM test sets, but it performed slightly better in predicting potentially dominant trees. DNNs were fitted in the python environment^73^ (version 3.8.5) using the PyTorch^74^ library (version 1.7.1). We constrained the training of the high-resolution DNN to 50 epochs, and thus a not fully plateauing cost function.

##### Cost functions

We used two cost functions to train low-resolution DNNs and one cost function to train the high-resolution DNN. As a baseline, we used the cross-entropy loss. The cross-entropy loss^24^ (CEL) function is a standard cost function for multiclassification with deep neural networks. It gets smaller the closer to zero the predicted probabilities for unobserved species are, and the closer to one the probability for the observed species is, and all species are assumed to be independent from each other. As a second cost function, we used the normalized discounted cumulative gain (NDCG). It evaluates model predictions up to a defined rank, which here we set to 500. If the observed species is present in the top 500 ranks, the model is rewarded, otherwise not. The reward thereby exponentially decreases, the higher the rank of the observed species within the predictions is. We implemented the NDCG with the LambdaNDCGLoss1 function of the pytorchltr module^75^, version 0.2.1.

##### Species distribution models

Single-species SDMs^11^ were built by relating occurrences and pseudoabsences to a subset of predictors selected from the low-resolution set. Three replicates of pseudoabsences (10’000) were generated with the geographic-specific approach and a background of environmentally stratified pseudo-absences as described in ref. ^15^. Species observations were thinned so that no more than one observation per 100×100 m grid cell was considered. Each species was modeled following an ensemble approach of different statistical techniques^76^, using five model algorithms with specific parameterizations: generalized linear models^77^ (GLMs) were implemented with second-order polynomials and logit link function; generalized additive models^78^ were implemented with thin-plate regression splines with maximum of four degrees of freedom, and logit link function; gradient boosting machines^79, 80^ were implemented with 5000 trees, an interaction depth of two, a learning rate of 0.001, and a Bernoulli distribution; random forests^81^ were implemented with 5000 trees and a minimum node size of one; and maximum entropy models^82^ were implemented with all feature types. For all algorithms, the total weights of occurrences and pseudoabsences were set to be equal in model calibrations and models were replicated three times with the different sets of pseudoabsences^83–85^. Ensemble predictions were derived by averaging the probabilistic predictions of all combinations of the five algorithms times three replicates. Species distribution modelling was conducted in the R environment, using the packages mgcv^86^, randomForest^87^, gbm^88^, and dismo^89^.

For each taxon, we used a variable-selection procedure to choose the best predictors. We tested all feasible combinations of predictors by running GLMs with second-degree polynomials and equal sums of weights for occurrences and pseudoabsences, and ranked them based on adjusted explained deviance^11^. Feasible combinations of predictors thereby were defined based on two criteria. Firstly, all absolute pairwise Pearson correlations within a predictor set had to be lower thanD0.7, so that collinearity was contained within the tolerable range^51, 90^. Secondly, the number of predictors was fixed for each taxon, and varied between three and ten as a function of the number of occurrences in the calibration dataset (26-35: three predictors; 36-45: four predictors; 46-55: five predictors; 56-65: six predictors; 66-75: seven predictors, 76-85: eight predictors, 86-95: nine predictors; and >95: ten predictors). This second criterion was implemented to achieve an approximate presence-to-predictor ratio ≥ 10^91, 92^. Taxa with 8-25 occurrences were modeled using ensembles of small models^93^, using the three best predictors selected from the variable selection procedure. Taxa with less than eight occurrences were not modeled.

For species distribution modeling, we used the low-resolution training data set as described above, but the taxonomic resolution considered was different. Models were fitted at the level of subspecies and species, and observations identified at the aggregate level were discarded. Habitat suitability projections at the aggregate level were created after model fitting, by taking weighted averages of the projections of all the species and subspecies that were part of the aggregate. Similarly, we combined projections at the subspecies and species level with weighted averages for species that were resolved to subspecies. The weights for these averages thereby were defined as the number of filtered presence observations of the corresponding taxon. Discarding aggregate-level observations for species distribution modelling led to a reduction of the data set that was small compared to the number of observations without date information that were discarded for multispecies DNN modeling.

#### Predictive analysis

##### Assessment of overall performance

We validated the predictions of multispecies DNNs and stacked SDMs (SSDMs) on citizen-science observations and survey data. All trained model configurations were evaluated against the held-out test set of the Info Flora observations, comparing ranks of observed species in the predictions, and top-1 and top-5 accuracies. As we were interested in the performance of typical field observations, we calculated weighted means of these statistics, using taxon-wise numbers of training observations as weights. We validated all trained models against the Info Flora test set, considering both the weighted (after 50 training epochs) and the unweighted (after 100 training epochs) versions of multispecies DNNs. We compared the predictive performance for 2438 taxa for which multispecies DNN and SSDM predictions existed with five observations per taxon, resulting in 12’190 observations. Three of these were removed due to incomplete coverage with environmental data.

Multispecies DNNs with the full set of predictors and SSDMs were also evaluated against data from the BDM survey which was not used for training. To this end, we first generated probabilistic predictions for each cell in the taxon-by-site presence/absence matrix. For multispecies DNNs, we approximated habitat suitability for each taxon and site as the 90^th^ percentile of observation probability from April to September. Then, for each taxon individually (i.e., for each column), we calculated the area under the receiver operating curve^30^ (AUC), indicating the capacity of the models to discriminate between locations where species are present and locations where species are absent. This assessment was based on 1339 taxa for which SDM and DNN predictions existed, and which were present in the test data. Similarly, we calculated AUC for each site individually (i.e., for each row), indicating the capacity of the models to discriminate species present at a site from species absent at a site. Site-by-site AUC was calculated based on the predictions for all 2438 taxa for which multispecies DNN and SSDM predictions existed, including those that were never observed in the BDM survey.

##### Timing of seasonal maxima in observation probability

We summarized seasonal observation probabilities pixel-wise by their maxima. For each pixel, we first predicted observation probabilities of the selected species with the high-resolution DNN from March to September. Then, we smoothed the seasonal response curves with a running mean using the ‘AvgPool1d’ function of the torch module with a kernel size of 22 days. Next, we identified the modes of the smoothed response curves, and extracted the timing of the highest mode. Finally, we masked pixels with smoothed maxima below 0.01 and those for which no modes could be identified, as these pixels were assumed to be unsuitable for the species. For the nationwide maps, timings of peak observation probability were aggregated to 200 m spatial resolution by median.

We evaluated how well *t_p_max_* corresponded to the date of Info Flora observations for which observed individuals were reported to be in ‘full bloom’. For a fraction of Info Flora observations information on phenology state existed, most of which reporting ‘full bloom’ for the observed individuals. For the common forbs *Ajuga reptans*, *Aster bellidiastrum*, *Galium pumilum* aggr., *Listera ovata*, *Orchis mascula*, and *Taraxacum officinale* aggr. we filtered observations with phenology state based on the same criteria as high-resolution training observations and matched them with predictions of *t_p_max_* as described above. Then, we calculated the median difference between *t_p_max_* and observation date (bias), and Spearman rank correlation to measure how well the predicted spatial patterns of *t_p_max_*correspond to observed patterns.

##### Mapping potentially dominant tree species

In order to map potentially dominant tree species, we first identified pixels with land cover classes including trees and shrubs, i.e., ‘normal dense forests’, ‘forest stripes, edges’, ‘forest fresh cuts’, ‘devastated forests’, ‘open forests (on agricultural areas)’, ‘open forest (on unproductive areas)’, ‘brush forest’, ‘groves, hedges’, clusters of trees (on agricultural areas)’, ‘trees in unproductive areas’, ‘scrub vegetation’, and ‘unproductive grass and shrubs’, as defined by ref. ^67^. For each of these wooded pixels, we then predicted observation probabilities for the species distinguished by the Swiss forest vegetation database from February to November and averaged them. Then, we quantified taxonomic reporting bias for each species as the ratio between summed cover-abundance percentages across the surveys of the Swiss forest vegetation database and the corresponding summed observation probabilities, and corrected observation probabilities for all pixels and species with these bias estimates. Finally, for each pixel, we identified the potentially dominant species as the one with the highest bias-corrected observation probability. For the nationwide maps, potentially dominant species were aggregated to 200 m spatial resolution by mode. As a sensitivity analysis, we generated equivalent maps for the northern Ticino region based on the low-resolution NDCG DNN and SDMs (Extended Data Fig. 4).

We validated predicted potentially dominant tree taxa against observed ones based on 1867 plots from the National Forest Inventory. As the coordinates of the NFI sites fell exactly on the corners of four pixels of our prediction layer, we extracted reweighted observation probabilities of all adjacent pixels and averaged them, before identifying the potentially dominant taxon. We used top-1 accuracy to measure overall prediction success and F_1_ score^94^ to estimate performance for individual species.

##### Projecting observation probabilities into the future

In order to project observation probabilities to 2071-2100, we first estimated future climatic conditions. To this end, we deduced expected deviations in temperature and precipitation for 2085 from the CH2018 projections of the National Centre for Climate Services^95^. In the region of the evaluated gradient, these projections estimated an increase in temperatures of about 4.75 °C and 4.25 °C for summer and winter respectively, a decrease in summer precipitation of about 27.5%, and an increase in winter precipitation of about 22.5% for the year 2085 and the emission scenario RCP8.5. We updated the corresponding low-resolution variables with these changes, and used a univariate generalized additive model with thin-plate regression splines to estimate future frost change frequency as a function of future winter temperature. The other low-resolution predictors were assumed to remain constant. We then predicted observation probabilities of potentially dominant graminoid species under current and future conditions, for May first, June first, July first, and August first, and applied a taxonomic-reporting-bias correction as described above but using survey data from the dry meadows and pastures initiative. For plotting, we smoothed the bias-corrected curves of observation probability along the gradient, using local polynomial regression fitting^96^ (R function loess) with a span of 0.15. As a sensitivity analysis, we also mapped bias-corrected habitat suitability based on SDM ensembles along the gradient under current conditions. Unless otherwise noted, model predictions were made in Python^73^ (version 3.8.5), while subsequent analyses as well as mapping were done in the R environment (version 3.6.3), using the packages ROCR^97^, terra^71^, and mgcv^86^.

## Acknowledgements

We thank all the experts and citizen scientists who contributed plant observations to the Info Flora database, the Swiss Biodiversity Monitoring program, the Swiss forest vegetation database, and the dry meadows and pastures initiative, as well as Tobias Roth and for help with BDM data extraction. This study was supported by the Swiss Data Science Center (SDSC) grants no. c17-07 (SPEEDMIND) and c19-09 (COMECO) to NEZ, DZ, PD, DNK and PB. PB, NEZ & DNK acknowledge additional funding from the WSL-internal project PLAPP and DNK & NEZ acknowledge additional funding from the 2019-2020 BiodivERsA joint call for research proposal under the BiodivClim ERA-Net COFUND programme (project ‘FeedBaCks’) with the national funder Swiss National Science Foundation (grant no. 20BD21_193907).

## Extended Data Figures and Tables

**Extended Data Fig. 1.**
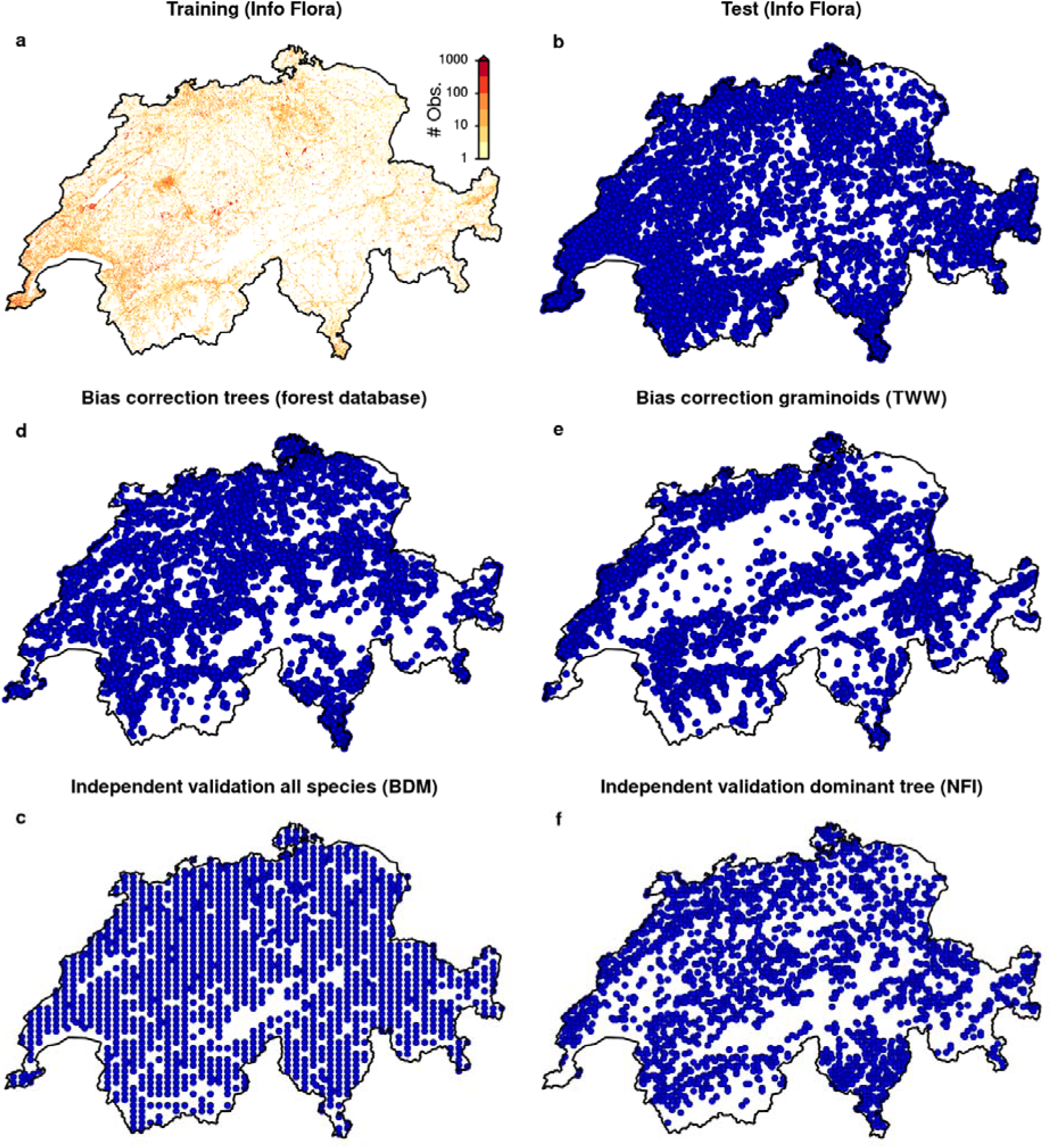
Spatial distributions of the observational data sets used in this study. **a**, number of quality-filtered training observations per 200×200 m pixel taken from the Info Flora database. **b**, locations of quality-filtered left-out test observations taken from the Info Flora database. **c**, locations of plant community observations for independent validation, taken from the Biodiversity Monitoring Program. **d**, locations of forest tree community observations for taxonomic bias correction, taken from the forest database. **e**, locations of grassland community plots for taxonomic bias correction, taken from the Dry Meadows and Pastures initiative. **f**, locations of forest tree community observations to validate model predictions taken from the National Forest Inventory.

**Extended Data Fig. 2.**
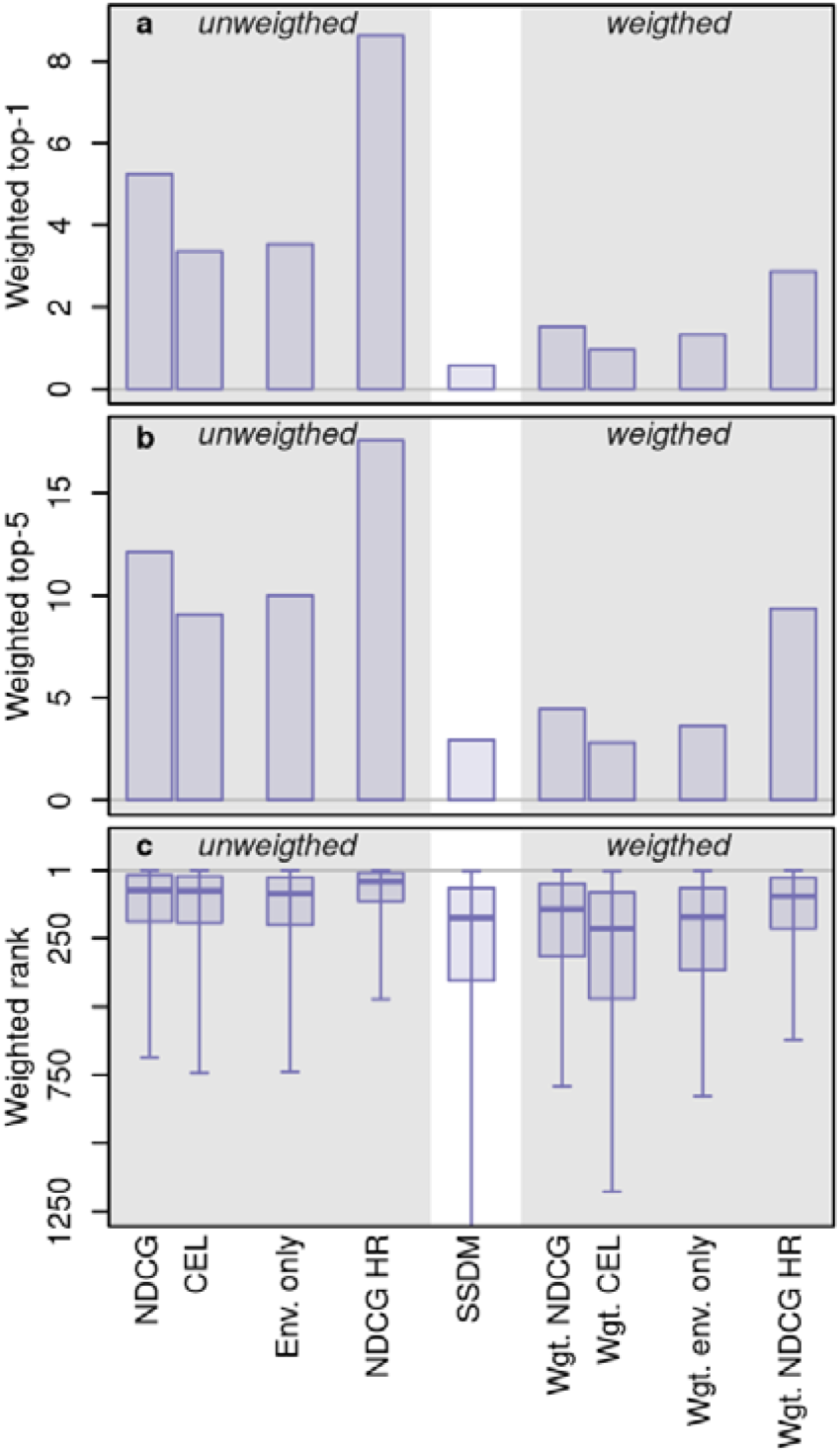
Overall performance of the trained deep neural networks and comparison to stacked species distribution models (SSDMs). Weighted versions were obtained by upsampling observations from rare taxa during training to obtain balanced taxon representation (see Methods). Top-1 accuracy in percent (**a**), top-5 accuracy in percent (**b**), and weighted ranks (**c**) are shown for various combinations of cost functions and predictor sets, as well as for predictions deduced from SSDMs. Central lines in the boxplots of panel (**c**) indicate weighted medians, boxes indicate weighted interquartile ranges, and whiskers indicate weighted 2.5 and 97.5 percentiles. Env. represents environmental predictors; NDCG represents the normalized discounted cumulative gain cost function; Wgt. represents weighted; CEL represents the cross-entropy loss cost function; and HR represents high resolution.

**Extended Data Fig. 3.**
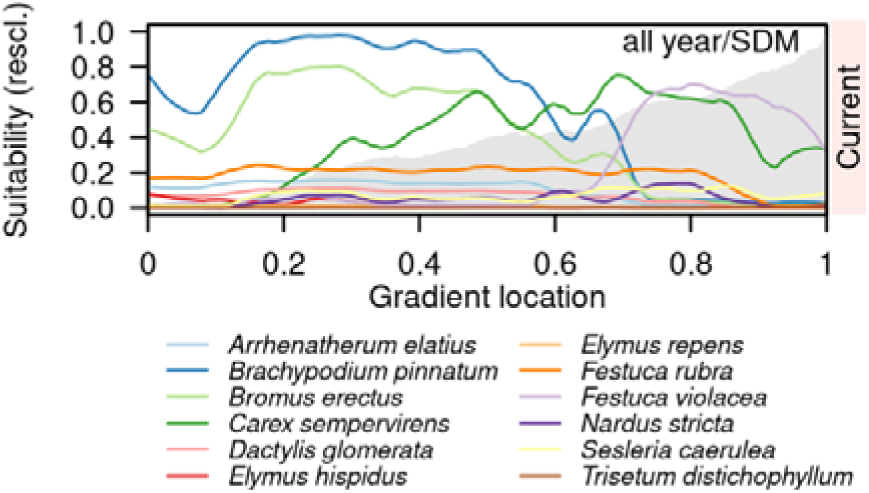
Reporting bias-corrected observation probabilities of graminoid taxa along an elevational gradient in the canton of Vaud (see Fig. 4). All-year habitat suitability under current conditions as derived from reporting-bias corrected predictions of stacked species distribution models. Gray shading in the background represents the elevation profile ranging from 400 to 2900 m asl. Illustrated are the twelve taxa with the most frequent potential dominance according to multispecies DNNs.

**Extended Data Fig. 4.**
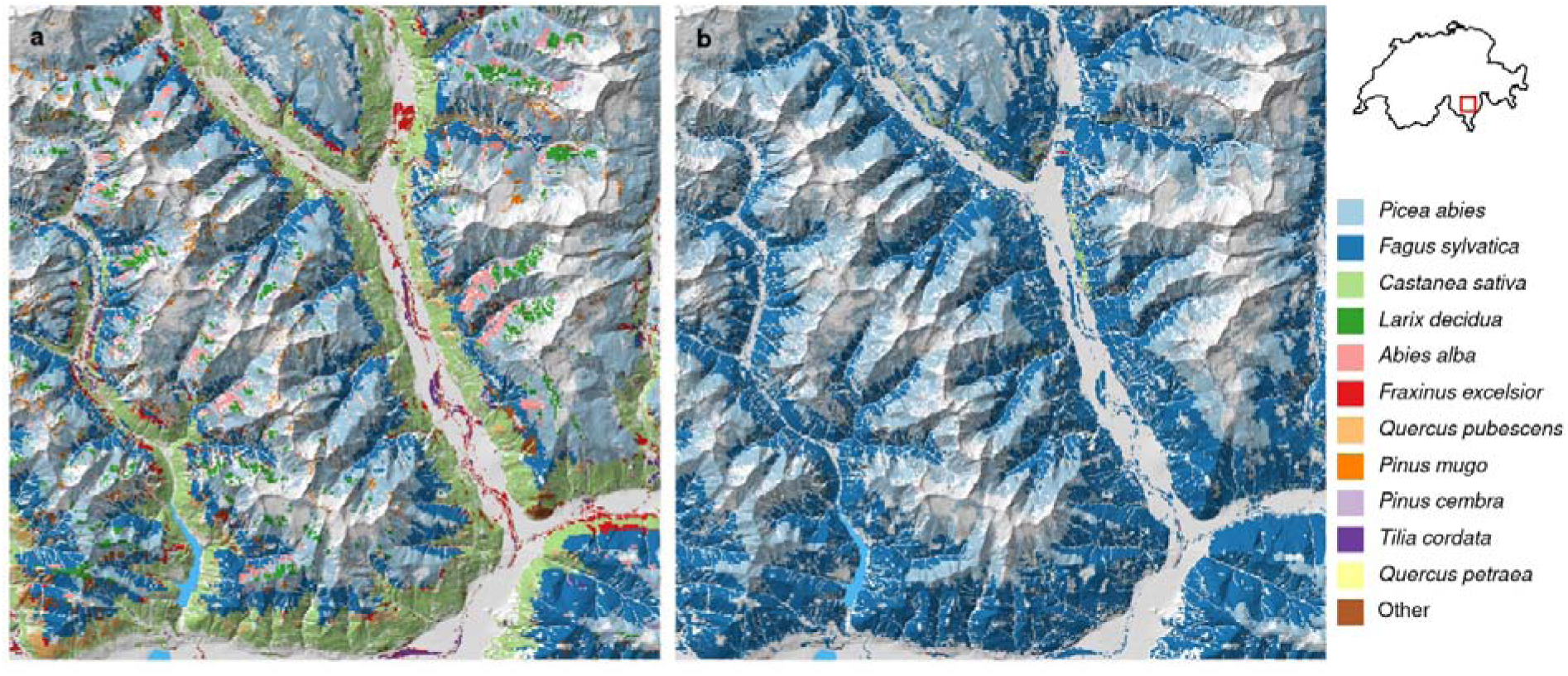
Potentially dominant canopy-forming tree species in wooded areas in Ticino. **a,** estimates according to reporting-bias corrected predictions of the low-resolution multispecies DNN with NDCG loss**; b,** estimates according to reporting-bias corrected predictions of stacked species distribution models. Colors represent species with the highest observation probability averaged from February to November (see legend). We compared observation probabilities of 37 tree species and tree species aggregates, as distinguished by the Walddatenbank (REF), and masked pixels with land cover classes with no expected trees (see methods).

**Extended Data Fig. 5.**
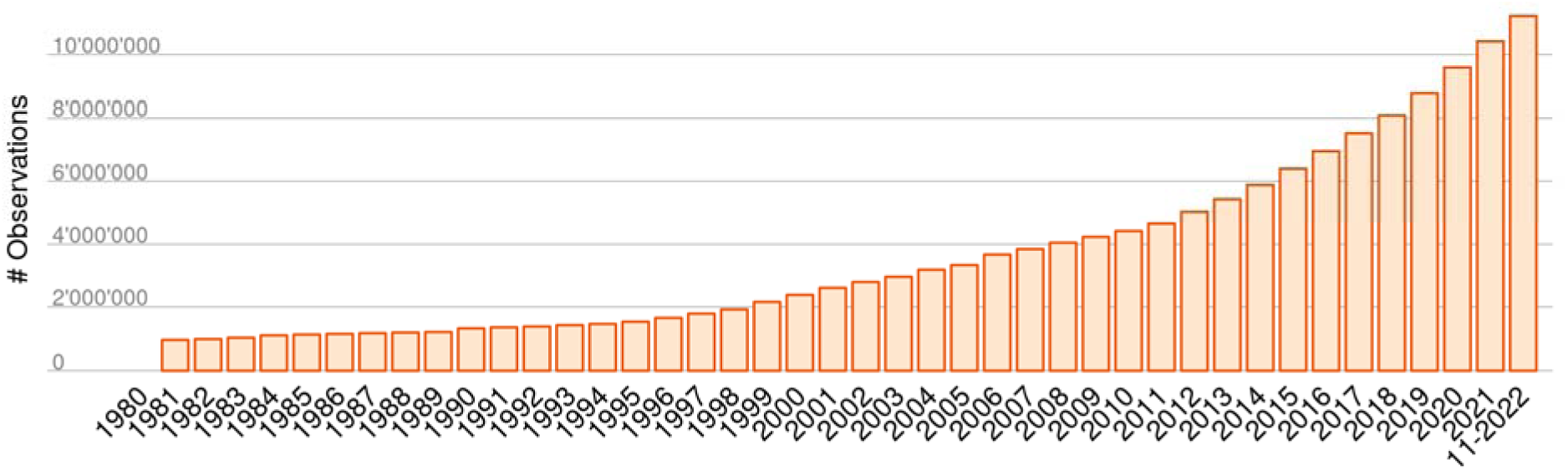
Increase of Info Flora observations for the period 1980-2019. Depicted are raw data before the filtering steps mentioned in the methods.

**Extended Data Fig. 6.**
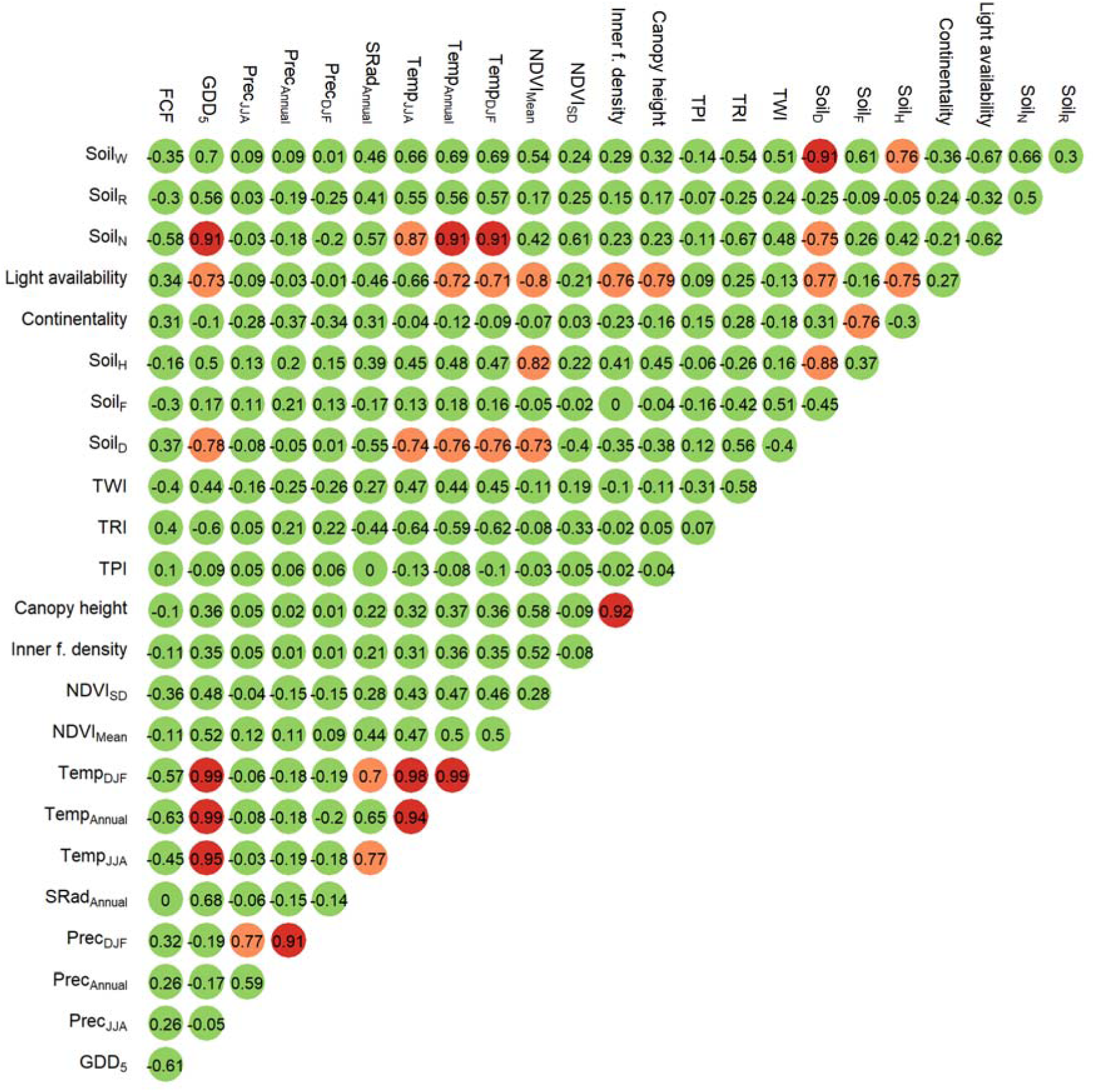
Pearson correlation coefficients between low-resolution environmental predictors across Switzerland. Green circles represent absolute Pearson correlation coefficients <0.7; orange circles represent absolute Pearson correlation coefficients between 0.7 and 0.9; and red circles represent absolute Pearson correlation coefficients higher than 0.9. Soil_W_, Soil_R_, Soil_N_, Soil_H_, Soil_F_, and Soil_D_, represent moisture variability, pH, nutrients, humus, moisture, and aeration; NDVI represents normalized difference vegetation index, Temp represents temperature, SRad represents solar radiation (direct and diffuse), Prec represents precipitation, GDD_5_ represents growing degree days above five degrees Celsius, and FCF represents frost change frequency. SD represents standard deviation, DJF represents December, January, February, and JJA represents June, July, August.

**Extended Data Table 1.** Selection of low-resolution environmental predictors. “Times selected in SDMs“ refers to the species-specific variable selection procedure described in the subsection “Species distribution models” of the methods and is a proxy for variable importance; “Correlation cluster #” gives a rough but imperfect indication of groups of predictors with absolute Pearson correlation coefficients > 0.7; “Selected” indicates the low-resolution predictors selected for DNNs (with “1”). Soil_W_, Soil_R_, Soil_N_, Soil_H_, Soil_F_, and Soil_D_, represent moisture variability, pH, nutrients, humus, moisture, and aeration; NDVI represents normalized difference vegetation index, Temp represents temperature, SRad represents solar radiation (direct and diffuse), Prec represents precipitation, GDD_5_ represents growing degree days above five degrees Celsius, and FCF represents frost change frequency. SD represents standard deviation, DJF represents December, January, February, and JJA represents June, July, August.

| Name | Times selected in SDMs | Correlation cluster # | Selected |
| --- | --- | --- | --- |
| TRI | 2224 | 10 | 1 |
| Soil<sub>R</sub> | 2103 | 13 | 1 |
| NDVI<sub>SD</sub> | 1794 | 7 | 1 |
| NDVI<sub>Mean</sub> | 1602 | 6 | 1 |
| SRad<sub>Annual</sub> | 1582 | 5 | 0 |
| Canopy height | 1577 | 8 | 1 |
| Prec<sub>JJA</sub> | 1496 | 4 | 1 |
| FCF | 1352 | 1 | 1 |
| Prec<sub>DJF</sub> | 1313 | 3 | 1 |
| Soil<sub>F</sub> | 1276 | 12 | 1 |
| TPI | 1185 | 9 | 0 |
| Soil<sub>N</sub> | 1117 | 2 | 0 |
| Soil<sub>W</sub> | 1056 | 2 | 0 |
| Continentality | 999 | 12 | 0 |
| TWI | 948 | 11 | 0 |
| Inner forest density | 642 | 8 | 0 |
| Temp<sub>JJA</sub> | 628 | 2 | 1 |
| GDD<sub>5</sub> | 524 | 2 | 0 |
| Soil<sub>D</sub> | 524 | 2 | 0 |
| Soil<sub>H</sub> | 501 | 6 | 0 |
| Prec<sub>Annual</sub> | 438 | 3 | 0 |
| Light availability | 412 | 6 | 0 |
| Temp<sub>Annual</sub> | 411 | 2 | 0 |
| Temp<sub>DJF</sub> | 365 | 2 | 0 |

**Extended Data Fig. 7.**
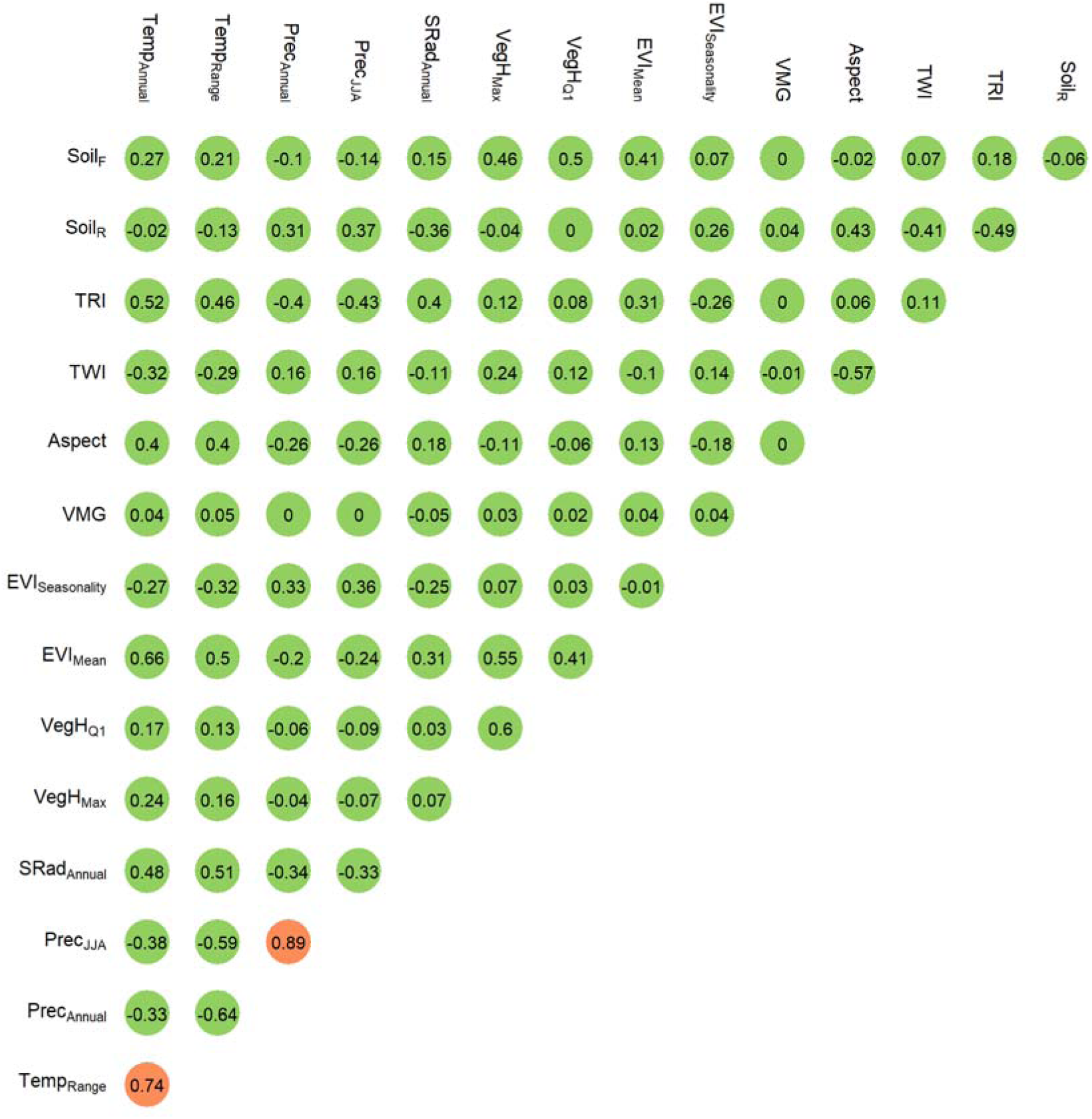
Pearson correlation coefficients between high-resolution environmental predictors across Switzerland. Green circles represent absolute Pearson correlation coefficients <0.7; orange circles represent absolute Pearson correlation coefficients between 0.7 and 0.9. Soil_R_ and Soil_F_ represent pH and nutrients, and respectively; EVI represents enhanced vegetation index, Temp represents temperature, Prec represents precipitation, SRad represents solar radiation (direct and diffuse), VegH represents vegetation height, and VMG represents forest canopy mixture (deciduous versus evergreen). Q1 represents first quartile, Max represents maximum, and JJA represents June, July, August

